# Upregulation of MAM by C99 disrupts ACSL4 activity and phospholipid homeostasis in Alzheimer’s disease models

**DOI:** 10.1101/2025.11.25.690197

**Authors:** J. Montesinos, T.D. Yun, I.D. Salomón-Cruz, S.C. Agudelo-Castrillón, M. Uceda, A.C. Ferre, C. Anton-Barros, N. Gomez-Lopez, R.R. Agrawal, D. Larrea, K.R. Velasco, A. Fernàndez-Bernal, E. Benitez, X. Zhu, E.A. Schon, F. Lopera, G.P. Cardona-Gomez, E. Area-Gomez

**Affiliations:** Department of Biomedicine, Centro de Investigaciones Biológicas Margarita Salas. CSIC, Madrid, Spain; Area of Cellular and Molecular Neurobiology, Group of Neurosciences, University of Antioquia, Medellín, Colombia; Institute of Human Nutrition, Columbia University Irving Medical Center, New York, NY, USA; Department of Neurology, Columbia University Irving Medical Center, New York, NY, USA; Metabolic Pathophysiology, Universitat de Lleida-Irblleida, Lleida, Spain; Universidad Europea de Madrid. Departamento de Ciencias Biomedicas. Madrid, Spain; Department of Pathology, Case Western Reserve University, School of Medicine, Cleveland, OH, USA; Department of Genetics and Development, Columbia University Medical Center, New York, NY, USA; Centro de Investigación Biomédica en Red sobre Enfermedades Neurodegenerativas (CIBERNED), Madrid, Spain

**Author notes:** Correspondence to: Dr. Area-Gomez, Department of Biomedicine, Centro de Investigaciones Biológicas Margarita Salas-CSIC, C/Ramiro de Maeztu, 9. Madrid (Spain). CIBERNED. Dr. Cardona-Gomez, Area of Cellular and Molecular Neurobiology, Group of Neurosciences, University of Antioquia, Medellín, Colombia, and Dr. Montesinos, Centro de Investigaciones Biológicas Margarita Salas-CSIC, C/Ramiro de Maeztu, 9. Madrid (Spain).,.

**Keywords:** Mitochondria-associated endoplasmic reticulum membranes (MAM), Alzheimer’s Disease, cholesterol, phospholipid, lipidomics, ACSL4

## Abstract

The structure and function of cellular and intracellular membranes are critically governed by the fatty acid (FA) composition of phospholipids (PLs), which is dynamically regulated by a network of enzymes that fine-tune lipid species according to cellular demands.

In this study, we identify a mechanism through which the formation of mitochondria-associated endoplasmic reticulum (ER) membranes (MAMs) modulates the activity of the acyl-CoA synthetase long-chain family member 4 (ACSL4), an enzyme that channels polyunsaturated fatty acids (PUFAs) into phosphatidylcholine (PC) via the Lands cycle. Through integrated biochemical, proteomic, and lipidomic analyses in both cellular and animal models, we demonstrate that MAM formation enhances ACSL4 activity, promoting arachidonic acid (AA) activation and its preferential incorporation into PC in concert with the MAM-localized lysophospholipid acyltransferase 4 (LPCAT4).

Our findings further uncover an unexpected link between this pathway and the pathogenesis of Alzheimer’s disease (AD). We show that elevated levels of C99—the β-secretase cleavage product of amyloid precursor protein (APP)—induce MAM remodeling through cholesterol clustering, which in turn activates ACSL4 and alters PC composition. This effect is mirrored in AD models as well as in fibroblasts, neurons, and immune cells derived from both familial and sporadic AD patients, all of which exhibit chronically increased C99 levels, heightened ACSL4 activity, and enrichment of PUFA-containing PC species, leading to lipid imbalance and membrane dysfunction.

Together, these results establish MAMs as dynamic lipid-regulatory hubs that coordinate ACSL4-dependent membrane remodeling and highlight the contribution of MAM dysregulation to lipid abnormalities observed in AD.

## Introduction

The lipid composition of cellular and intracellular membranes defines their characteristics, such as permeability and curvature, to facilitate processes such as endocytosis and exocytosis, as well as the function of integral proteins, including receptors and ion channels. Defects in the regulation of cellular lipid metabolism can influence biophysical properties of membranes and induce organellar and cellular dysfunction, particularly in those cell types with highly active and complex membranes, such as neurons [1, 2]. Not surprisingly, mutations in lipid regulation genes and disturbances in the lipidomes of various tissues are often associated with neurological disorders and neurodegenerative diseases such as Alzheimeŕs disease (AD) [3, 4].

A major determinant of membrane properties, such as fluidity and permeability, is the fatty acid (FA) composition of constituent phospholipids (PL) and the length and degree of FA saturation. Through this impact on membrane properties, FAs can modulate multiple cellular events [5, 6]. Additionally, FAs released from PLs can act as signaling molecules and as precursors of metabolites such as eicosanoids and hormones [7, 8]. The fatty acyl composition of cellular PLs is therefore under strict regulation, with the FA composition of phosphatidylcholine (PC), the most abundant PL in mammalian cells, being especially regulated.

During the *de novo* synthesis of PLs, PC and other PLs are derived from phosphatidic acid precursors; however, once synthesized, their FA composition is rapidly modified by deacylation and reacylation reactions via the Lands cycle based on cellular needs and environmental conditions [9]. In this pathway, PLs are deacylated by phospholipases and reacylated by lysophospholipid acyltransferases. In the case of PC, lysophosphatidylcholine acyltransferases (LPCATs) catalyze the insertion of specific FAs into PC, generally at the sn-2 position [10]. These FAs can either be synthesized *de novo* or derived from the diet and transported into the cell, where they are first activated by the addition of a coenzyme A group (CoA) by Acyl-CoA synthetases (ACSL1-6) [11]. Once activated, FA-CoA can then be modified by specific membrane-bound elongases (ELOVLs) and desaturases (FADs) [12, 13].

In mammals, the FA elongation process generates a two-carbon-unit chain extension by the elongase isoforms 1–7 (ELOVL1–7), each with distinct cell type-dependent expression and substrate preference, ELOVL1-4 and ELOVL6-7 have a preference for saturated and monounsaturated FAs while ELOVL5 has a preference for polyunsaturated FAs (PUFAs) [12]. In addition to undergoing elongation, FAs can also undergo desaturation catalyzed by three distinct desaturase enzymes - SCD1, FADS1 and FADS2 – carrying out Delta-9, Delta-5 and Delta-6-desaturase activities, respectively, to introduce double bonds at specific sites (omega [ω] sites) along the FA chain [13]. The coordinated activity of different isoforms of ACSLs, elongases and desaturases results in the orchestrated production and metabolism of FAs. These can be derived via *de novo* synthesis (ω-9 FA) or from dietary essential FA precursors (ω-3 and ω-6 FA) and can be converted into PUFAs, such as arachidonic acid (AA, C20:4n-6) and docosahexaenoic acids (DHA, C22:6n3), with important roles in maintaining membrane integrity and regulating cellular signaling events, especially in the CNS [5]. These newly synthesized FA-CoAs can then enter various metabolic pathways, including β-oxidation and phospholipid synthesis and reacylation, to maintain the composition and fluidity of cell membranes. In addition, FA-CoAs can function as signaling molecules during gene expression, cell division, differentiation, and other pathways, such as inflammation [14].

Owing to their pivotal role in the modulation of cellular membranes, the expression and activity of ACSLs, FADSs and ELOVLs is kept under a tight and complex regulation by multiple transcriptional factors (e.g. LXR, PPARg, SREBP1c) in a tissue- and cell type-specific manner, and in response to different environmental conditions [15]. In particular, the regulation of FA activation by ACSLs has been shown to be essential not only for FA synthesis but also for channeling FA into different pathways [15].

It is noteworthy that different ACSL isoforms have preferences for different FAs. For example, ACSL4-6 have a marked preference for several PUFAs, such as AA, DHA and eicosapentaenoic acid (EPA), whereas ACSL1 prefers saturated and monounsaturated FAs that are 16–18 carbons in length [7, 11]. In addition to their FA preferences, the capacity of different ACSLs to direct FA-CoAs towards oxidation or to the synthesis of other lipids is regulated by their subcellular localization and association with different membrane domains [16]. This compartmentalization of ACSLs allows for specific FA-CoA species to be handed off to lipid biosynthetic and oxidative enzymes while appropriately limiting their access to other pools [17]. Notably, ACSL4 (also called fatty acid CoA ligase 4, FACL4), an isoform highly expressed in brain tissues, has been shown to localized to mitochondria-associated membranes (MAM) within the endoplasmic reticulum (ER); however, the functional significance of this localization has not yet been determined [18].

MAM is a dynamic lipid-raft domain within the ER that transiently clusters and regulates the activity of multiple metabolic enzymes in response to cellular needs. Specifically, the formation of the MAM has been shown to result in the association and activation of various enzymes involved in the synthesis of phospholipids, sphingolipids and neutral lipids [19], along other important metabolic functions such as calcium transfer and regulation of mitochondrial dynamics. Importantly, the formation of MAM domains and the activities regulated by this domain are upregulated in AD cells. We previously showed that the accumulation of the APP-derived C99 fragment at MAM leads to the buildup of cholesterol at MAM, thus upregulating MAM-located activities [20]. However, the consequences and significance of MAM defects in the cell are still under investigation.

We show here that MAM formation driven by C99 induces ACSL4 activity and the reacylation of PC with AA. Moreover, our data indicate that the upregulation of this mechanism in cell and animal models of AD, as well as in fibroblasts from familial and sporadic AD cases, contributes to the lipid abnormalities frequently observed in AD. Our data indicate that the regulation of MAM is essential in the maintainance of phospholipid saturation and thus, the composition and function of cellular membranes.

## Results

### ACSL4 is enriched and activated in MAM domains

Previous studies have shown that ACSL4, a long-chain acyl-CoA synthetase highly expressed in brain tissue, is localized predominantly to mitochondria-associated membranes (MAMs) within the ER [21]. However, the functional significance of this localization in regard to ACSL4 enzymatic activity remains unexplored. To address this, we first quantified ACSL4 protein levels in subcellular fractions isolated from mouse brain tissues (**Fig. 1A**). As reported previously [22], although ACSL4 is present in the bulk ER, it is significantly enriched in purified MAM fractions.

**Figure 1:**
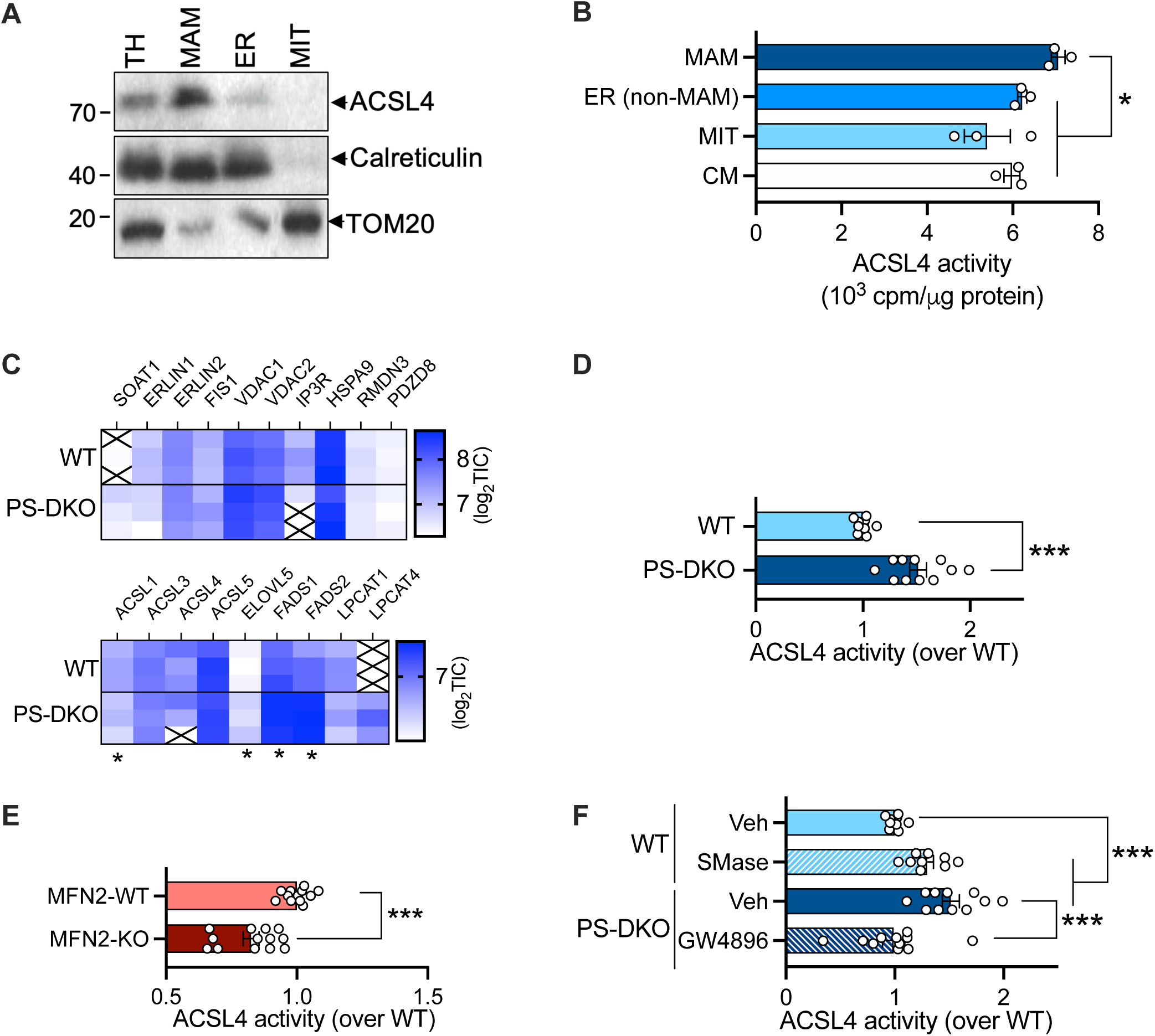
Regulation of ACSL4 activity at MAM domains. **A.** Representative western blot of ACSL4 levels in mouse brain. Calreticulin and TOM20 were used as loading controls for ER and mitochondria (MIT), respectively. TH: total homogenate. **B.** ACSL4 activity in subcellular fractions obtained from C57BL6 mouse brain. CM: crude membranes. MIT: mitochondria. ER: Endoplasmic Reticulum (non-MAM “bulk” fraction). **C.** Heatmap of fold changes of the log_2_(Total Ion Current) from PhotoClick-cholesterol pulldowns of MAM obtained from WT or PS-DKO cells. N=3. **D.** ACSL4 activity measured from membrane preparations of WT or PS-DKO MEFs. **E.** ACSL4 activity measured from membrane preparations of MFN2-WT or MFN2-KO MEFs. **F.** ACSL4 activity measured from membrane preparations of WT or PS-DKO cells treated with sphingomyelinase (SMase) or the SMase inhibitor GW4896, respectively.

ACSL4 preferentially activates long-chain PUFAs, with a particular affinity for Δ5 fatty acids, such as AA [8]. We assessed ACSL4 activity by measuring the rate of acylation of exogenously-added ^3^H-arachidonic acid (^3^H-AA) in each isolated subcellular fraction and found that ACSL4 activity was significantly higher in the MAM fraction than in other compartments (**Fig. 1B**).

MAMs are dynamic structures that transiently cluster and regulate multiple metabolic enzymes [19]. To validate the presence of ACSL4 in MAMs, we performed proteomic profiling in MAMs isolated from mouse embryonic fibroblasts (MEFs) lacking presenilin-1 and -2 (*PSEN1/2* double knockout, PS-DKO), a model where MAMs are constitutively upregulated, and MAMs isolated from wild-type (WT) MEFs [23]. Cells were incubated with a PhotoClick-cholesterol analog that integrates into cellular membranes [20, 24]. Upon UV crosslinking, the analog covalently binds any interacting proteins. After MAM isolation, PhotoClick-cholesterol-interacting proteins were tagged with biotin via click chemistry, enriched through streptavidin pull-down, and analyzed by LC-MS/MS (**Fig. 1C**).

This unbiased proteomics analysis identified 1647 proteins in at least 2 out of the 3 replicates per condition (**Table S1**), including ACSL4, as expected. In this dataset, we noted the enrichment of ER and mitochondrial proteins (**Fig. S1A & Table S2**) and classical markers of MAM such as SOAT1, ERLIN1/2 and tethers of ER-mitochondria contact sites such as FIS1/BAP31 [25], VDAC/IP3R/GRP75 [26], RMDN3 (also called PTPIP51) [27], and PDZD8 [28]. When comparing MAM proteomes from WT and PS-DKO samples, 94 proteins were exclusively present in all replicates obtained from WT cells whereas 28 proteins were exclusively present in PS-DKO replicates. 119 proteins were present in all WT replicates but in 1 PS-DKO replicate, while 32 proteins were present in converse conditions (“semiexclusive hits”, **Table S3**). Differential enrichment comparison revealed that 110 hits were significantly altered between groups (**Table S4**). Specifically, 41 proteins were significantly enriched while 69 proteins were depleted in the MAM fractions of PS-DKO cells when compared to WT samples (**Fig. S1B & Table S5**).

Analysis of the data set obtained within KEGG classes by Gene Ontology revealed a significant enrichment of 49 important cellular categories (**Table S6**). Specifically, several categories pointed to FA metabolic pathways: ‘fatty acid metabolism’, ‘fatty acid degradation’, ‘biosynthesis of unsaturated fatty acids’ and ‘fatty acid elongation’ (**Fig. S1C**). In this way, we confirmed the enrichment of ACSL4 in MAM fractions, along with other FA metabolism-related enzymes, such as fatty acid desaturases 1 and 2 (FADS1/2), lysophosphatidylcholine acyltransferase isoforms 1, 3 and 4 (LPCAT1/3/4), and ELOVL5. Notably, in MAM fractions from PS-DKO cells, levels of FADS1, FADS2, and LPCAT4 were significantly increased, while ACSL4 levels remained unchanged compared to wild-type controls (**Fig. 1C & Fig. S1C**).

These findings were corroborated by western blot analyses, which revealed increased protein levels of FADS1/2 in PS-DKO MAMs, with no significant changes in ACSL4 abundance (**Fig. S1D**). Despite this, MAM fractions from PS-DKO cells displayed significantly higher ^3^H-AA incorporation, implying elevated ACSL4 enzymatic activity (**Fig. 1D**). This result supports previous observations [22], suggesting that ACSL4 activity is modulated independently of its expression level, but rather by its subcellular localization. In agreement, cells deficient in the formation of MAM due to the ablation of Mitofusin-2 (MFN2), a protein critical for ER–mitochondria tethering [22], exhibited significant reductions in ACSL4 activity (**Fig. 1E**) with no effect on ACSL4 levels (**Fig. S1E**).

The formation of MAMs can be stimulated by increased ER cholesterol levels via transfer from the plasma membrane (PM) [29]. This process can be induced experimentally by treating cells with recombinant sphingomyelinase (SMase), which hydrolyzes sphingomyelin at the PM [30]. Accordingly, to test further whether MAM formation regulates ACSL4 activity, we treated WT MEFs with SMase and measured ^3^H-AA activation. SMase treatment led to a significant increase in ACSL4 activity (**Fig. 1F**). Conversely, treatment with the SMase inhibitor GW4896, which prevents MAM formation [20, 31], restored elevated ACSL4 activity in PS-DKO cells to wild-type levels (**Fig. 1E**). Neither of these treatments altered ACSL4 total protein levels (**Fig. S1F**).

Taken together, our results agree with previous evidence [22], indicating a discordance between protein abundance and ACSL4 enzyme activity, and that the latter is mainly regulated by the formation of MAM domains.

### Upregulation of MAM promotes arachidonic acid incorporation into phosphatidylcholine

ACSL4 is known to preferentially activate PUFAs, particularly AA, for phospholipid (PL) synthesis and remodeling, rather than for β-oxidation or triglyceride (TAG) synthesis [8]. To investigate how MAM upregulation and ACSL4 activation affects fatty acid (FA) routing into different lipid classes, we incubated PS-DKO and WT cells with radiolabeled saturated palmitic acid (^3^H-PA), monounsaturated oleic acid (^3^H-OA), or polyunsaturated arachidonic acid (^3^H-AA), and analyzed the incorporation of these FAs into lipid species using thin-layer chromatography (TLC) (**Fig. 2A-B**).

**Figure 2:**
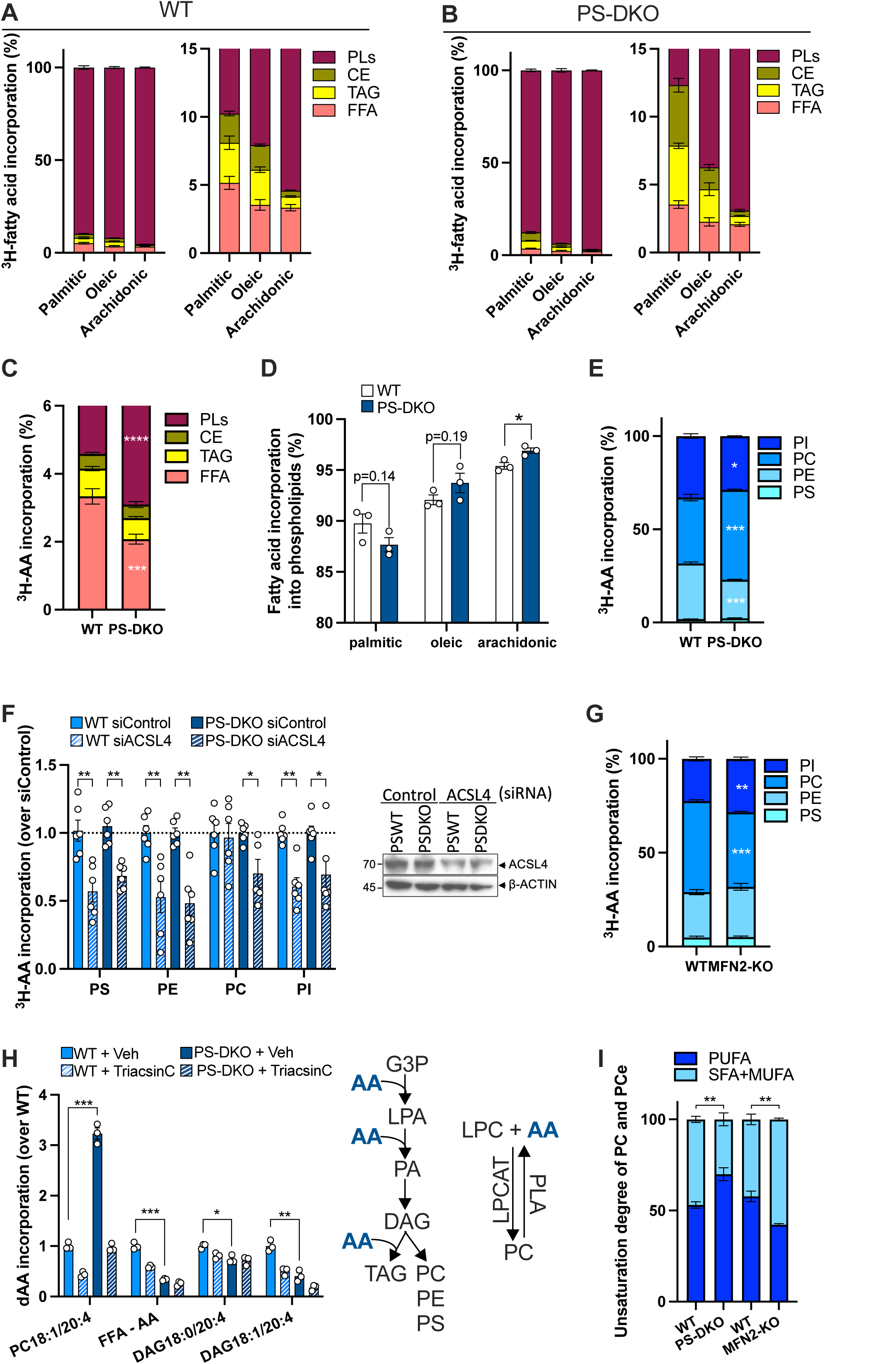
Arachidonic acid incorporation into phosphatidylcholine is mediated by ACSL4. A, B, **C.** Palmitic, oleic or arachidonic acid incorporation into phospholipids (PLs), cholesteryl esters (CE), and triacylglycerides (TAG) was measured upon 4h incubation in **(A)** WT and **(B)** PS-DKO cells. Free fatty acid (FFA) levels were also measured. A comparison of arachidonic incorporation in WT and PS-DKO cells is shown in **(C)**. Fig. S2A shows the comparison for palmitic and oleic acid. N=3. **D, E.** Incorporation of palmitic, oleic or arachidonic acid into phosphatidylserine (PS), phosphatidylethanolamine (PE), phosphatidylcholine (PC) and phosphatidylinositol (PI) was analyzed upon 4h incubation. A comparison of arachidonic acid incorporation into specific phospholipid classes in WT and PS-DKO cells is shown in **(E)**. Fig. S2B shows the comparison for palmitic and oleic acid. **F.** Incorporation of ^3^H-arachidonic acid upon 4h incubation into specific phospholipid species was assessed in WT and PS-DKO cells silenced for ACSL4. A representative western blot of ACSL4 is included to show the efficiency of the silencing (∼70%). Incorporation into TAG and CE is shown in Fig. S2C. **G.** Incorporation of arachidonic acid into specific phospholipid classes was assessed in WT and MFN2-KO cells. N=3. **H.** WT and PS-DKO cells were incubated for 4h with deuterated arachidonic acid (dAA) in the presence of the ACSL inhibitor Triacsin C and dAA incorporation was analyzed by lipidomics. Incorporation into phosphatidylcholine (PC) and diacylglycerol (DAG) is shown. A complete heatmap of dAA incorporation is shown in Fig. S2F. Left diagram depicts the *de novo* synthesis of phospholipids (PC, PE, PS) from glycerol-3-phosphate (G3P). A fatty acid is first added to G3P to form lysophosphatidic acid (LPA). A second fatty acid is then added to phosphatidic acid (PA). PA is subsequently dephosphorylated into DAG which can serve as a substrate for the synthesis of either TAG or phospholipids. Right diagram shows the Lands cycle that facilitates the exchange of fatty acids within phospholipids. This cycle begins with the removal of a fatty acid from PC, catalyzed by phospholipase A (PLA), which creates lysophosphatidylcholine (LPC). LPC can then be reacylated with a new fatty acid by LPCAT. **I.** The degree of unsaturation, calculated as the percentage of SFA+MUFA (presenting 0 or 1 degree of unsaturation) vs PUFA (presenting more than 1 degree of unsaturation), of PC and its ether-linked form (PCe) were analyzed in PS-DKO cells, MFN2-KO cells and their respective WT controls. N=3.

As expected, all exogenous FAs were incorporated predominantly into PLs among the lipid classes assessed. However, PS-DKO cells showed a significant increase in ^3^H-AA incorporation into PLs, along with a corresponding decrease in the unesterified pool of free ^3^H-AA forms, compared to control cells (**Fig. 2C-D**). Interestingly, PS-DKO cells also exhibited increased channeling of ^3^H-PA into cholesteryl esters (CEs), whereas ^3^H-OA incorporation remained unchanged across lipid classes, compared to controls (**Fig. S1A**). Further analysis of ^3^H-AA incorporation into specific PL subclasses (**Fig. 2E & Fig. S2B**) revealed that, in WT cells, AA was fairly evenly distributed among phosphatidylethanolamine (PE), phosphatidylcholine (PC), and phosphatidylinositol (PI), with little incorporation into phosphatidylserine (PS). In contrast, PS-DKO cells showed a preferential incorporation of ^3^H-AA into PC compared to other PL classes.

To validate the role of ACSL4 in the differential routing of AA, we repeated the labeling experiments in both control and PS-DKO cells following ACSL4 silencing by siRNA. While incorporation of ^3^H-AA into CE and TAG was unaltered (**Fig. S2C**), knockdown of ACSL4 reduced ^3^H-AA incorporation into PS, PE, and PI in both genotypes (**Fig. 2F**). However, in control cells, ACSL4 knockdown had little effect on AA incorporation into PC, whereas in PS-DKO cells, silencing ACSL4 significantly reduced ^3^H-AA incorporation into PC. These results suggest that, under basal conditions, AA is not a preferred substrate for PC synthesis via *de novo* Kennedy pathway. However, in cells with constitutively active MAMs, elevated ACSL4 activity redirected AA-CoA mainly into the PC synthesis pathway. Supporting this, MFN2 knockout (MFN2-KO) cells, which have impaired MAM formation and reduced ACSL4 activity, showed the opposite phenotype (**Fig. 2G & Fig. S2D**).

The remodeling of PC is mediated primarily through the reacylation of existing PC species via the Lands cycle, rather than during the PC synthesis [32]. In this process, phospholipase A2 (PLA2) hydrolyzes the fatty acid at the sn-2 position of PC, producing lysophosphatidylcholine (LPC). Subsequently, LPC acyltransferase (LPCAT) enzymes re-esterify the LPC with additional FA [9]. Notably, LPCAT isoforms 3 and 4 (LPCAT3 and LPCAT4) demonstrate a strong preference for utilizing activated AA and other PUFAs to acylate the sn-2 position of PC [32]. To determine the contribution of PC reacylation via the Lands cycle to the incorporation of AA into PC in our cell models, we treated cells with methyl arachidonyl fluorophosphonate (MAFP), an irreversible inhibitor of PLA2 activity. Interestingly, inhibition of PLA2 led to higher levels of ^3^H-AA incorporation into PC in PS-DKO cells compared to controls, suggesting increased LPCAT activity in the mutant cells under conditions of MAM upregulation (**Fig. S2E**).

To further validate this finding, we monitored and quantified the incorporation of deuterated AA (dAA) into various phospholipid species in our cell models using lipidomics analysis (**Fig. 2H & Fig. S2F**). As a control, we treated the cells with Triacsin C, which inhibits ACSL4 [33]. Under these conditions, dAA in WT cells was incorporated primarily into specific species of PE and plasmalogen PE (PEp). Exogenous dAA was also detected in the corresponding diacylglycerol (DAG) precursors of these phospholipids in control cells, indicating that dAA-containing PE species were mainly synthesized *de novo*. Conversely, PS-DKO cells exhibited a clear preference for incorporating dAA into PC (notably PC 18:1/20:4), and to a lesser extent into PS, compared to controls. However, we did not observe a corresponding increase in dAA incorporation into DAG species. This suggests that, unlike in WT cells, dAA incorporation into PC in PS-DKO cells occurs primarily through LPCAT-mediated reacylation, consistent with enhanced activity of the Lands cycle.

We next asked whether these changes in ACSL4-mediated FA incorporation led to alterations in cellular lipid composition of our cell models. To test this, we analyzed and quantified PC molecular species in cells with either upregulated (PS-DKO) or disrupted (MFN2-KO) MAMs by lipidomics (**Fig. 2I & S2G**). PS-DKO cells displayed a significant enrichment of polyunsaturated PC species and their ether-linked analogs (PCe), whereas MFN2-KO cells exhibited a reduction in these same species. Importantly, ACSL4 knockdown significantly reversed this increase in PC-PUFA levels in PS-DKO cells, compared to controls (**Fig. S2H**).

Taken together, these findings support the model that MAM formation enhances ACSL4 activity, leading to the preferential channeling of AA-CoA into PC. This process substantially remodels the cellular lipidome, particularly enriching PC species with PUFAs.

### The C99-cholesterol interaction drives MAM remodeling and ACSL4 activation

We previously demonstrated that C99, the β-secretase cleavage product of APP, can induce MAM formation by clustering cholesterol within the ER [20]. To determine whether the physical interaction between C99 and cholesterol is sufficient to activate ACSL4, we compared the effects of wild-type C99 (C99^WT^) with a cholesterol-binding-deficient mutant (C99^MUT^). The C99^MUT^ variant harbors alanine substitutions (G700A, I703A, G704A), which significantly reduce its affinity for cholesterol [20]. We transiently expressed either C99^WT^ or C99 ^MUT^ in APP/APLP2 double knockout (APP-DKO) MEFs, which lack endogenous *APP* and its homolog *APLP2*. Cells transfected with an empty vector (EV) served as controls. Twenty-four hours post-transfection, cells were treated for 16 hours with the γ-secretase inhibitor DAPT to block further processing of C99, allowing accumulation of the full-length C99 protein. Strikingly, expression of C99^WT^ led to significant upregulation of ACSL4 activity, which was not observed in cells expressing C99^MUT^, indicating that C99’s ability to bind or cluster cholesterol is essential for promoting ACSL4 enzymatic activity (**Fig. 3A**).

**Figure 3:**
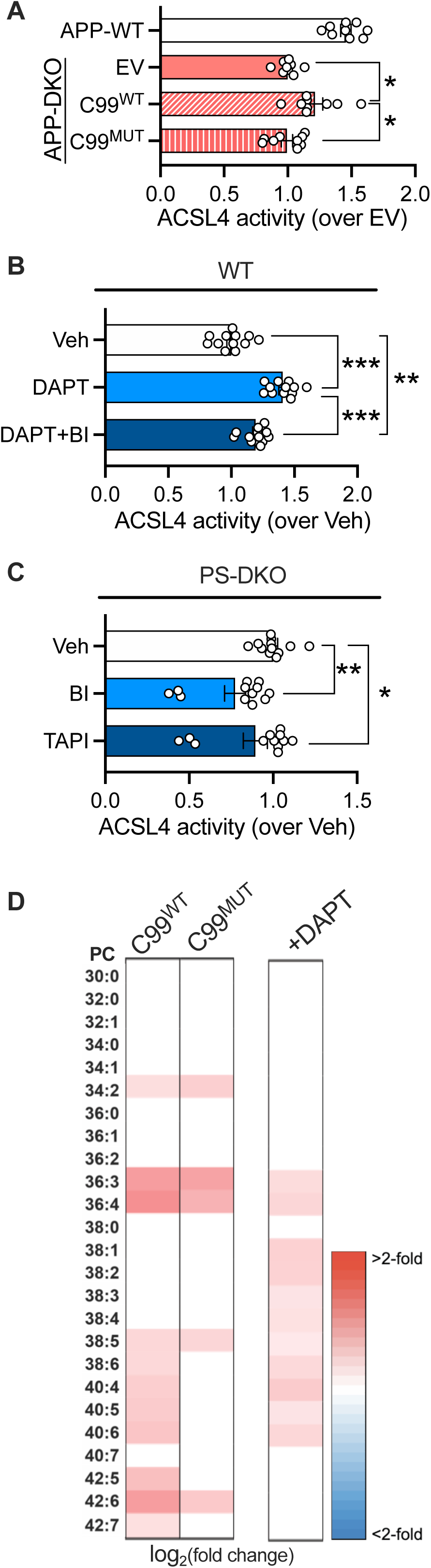
The C99-cholesterol interaction drives upregulation of ACSL4 activity and lipidomic remodeling. **A.** ACSL4 activity measured from membrane preparations of *APP/APLP2* double knock-out (APP-DKO) cells overexpressing C99^WT^ or its cholesterol binding-deficient version (C99^MUT^). An empty vector (EV) and APP-WT cells were used as controls. **B.** ACSL4 activity was assayed in WT MEFs treated for 16h with DAPT or a combination of DAPT and BACE1 inhibitor (BI). DMSO treatment was used as vehicle (VEH). **C.** PS-DKO MEFs were treated for 16h with BI or an α-secretase inhibitor (TAPI) and ACSL4 activity was measured. DMSO treatment was used as vehicle (VEH). **D.** Heatmap of fold changes of the levels of PC species analyzed by lipidomics from APP-DKO MEFs transfected with C99^WT^ or C99^MUT^ over the EV control (left columns). Heatmap of PC species of DAPT-treated WT MEFs over vehicle-treated control (right column). Only significant changes are coloured. N=4-5.

Pharmacological inhibition of γ-secretase with DAPT, known to promote C99 accumulation and enhance MAM activity [31], also resulted in increased ACSL4 activity (**Fig. 3B**). Conversely, inhibition of β-secretase (BI), which prevents C99 formation, led to decreased ACSL4 activity in DAPT-treated WT cells and in PS-DKO cells (**Fig. 3B-C**). Consistent with findings in PS-DKO cells, increased C99^WT^ expression and the accompanying upregulation of MAM was associated with elevated levels of PUFA-containing phosphatidylcholine (PC-PUFA) species in the lipidome (**Fig. 3D**). In contrast, this enrichment was attenuated in cells expressing the C99^MUT^ construct, highlighting the requeriment of C99 cholesterol-binding for the modulation of lipid composition. On the other hand, the levels of ether-linked PC (PCe) species remained unchanged under these conditions.

In conclusion, our data indicate that the activation of MAM by the accumulation of C99 can induce significant alterations in PC homeostasis via the upregulation of ACSL4.

### ACSL4 activity is upregulated in Alzheimer’s disease models

C99 levels and MAM activity are increased in tissues and cellular models of Alzheimer’s disease (AD), including those from mice expressing mutant forms of presenilin-1 (PS1-KI^M146V^ knock-in mice) or APP and Tau (3xTg-AD mouse model: APP^Swedish^, PS1^M146V^, Tau^P301L^), as well as in cells derived from both early- and late-onset AD patients. To further explore the relationship between MAM upregulation and ACSL4, we evaluated ACSL4 protein levels in immature brain tissue (P0) of PSEN1^M146V^-KI and in 3xTg-AD mice in comparison to C57BL6 wild-type mice. A slight tendency to reduce ACSL4 was observed (**Fig. S3A**). This change was supported by a decrease of ACSL4 immunofluorescence detection in mature neurons (NeuN^+^), both in PSEN1^M146V^-KI and 3xTg-AD mice (**Fig. S3B**). Next, we assessed ACSL4 enzymatic function in primary neurons derived from neonatal PS1^M146V^-KI mice (**Fig. 4A**) and in aged (1 year-old) brain homogenates (**Fig. 4B**) from the same model. In both primary neuronal cultures and brain tissue, ACSL4 activity was significantly increased compared to wild-type controls. As before, this elevation was recapitulated in wild-type neurons treated with a γ-secretase inhibitor, which promotes the accumulation of C99 and enhances MAM formation. Conversely, treatment of PS1^M146V^-KI neurons with a β-secretase inhibitor, which prevents C99 generation, restored ACSL4 activity to wild-type levels (**Fig. 4C**). These results suggest that C99 acts as an upstream regulator of ACSL4 activity during early stages of cellular dysfunction.

**Figure 4:**
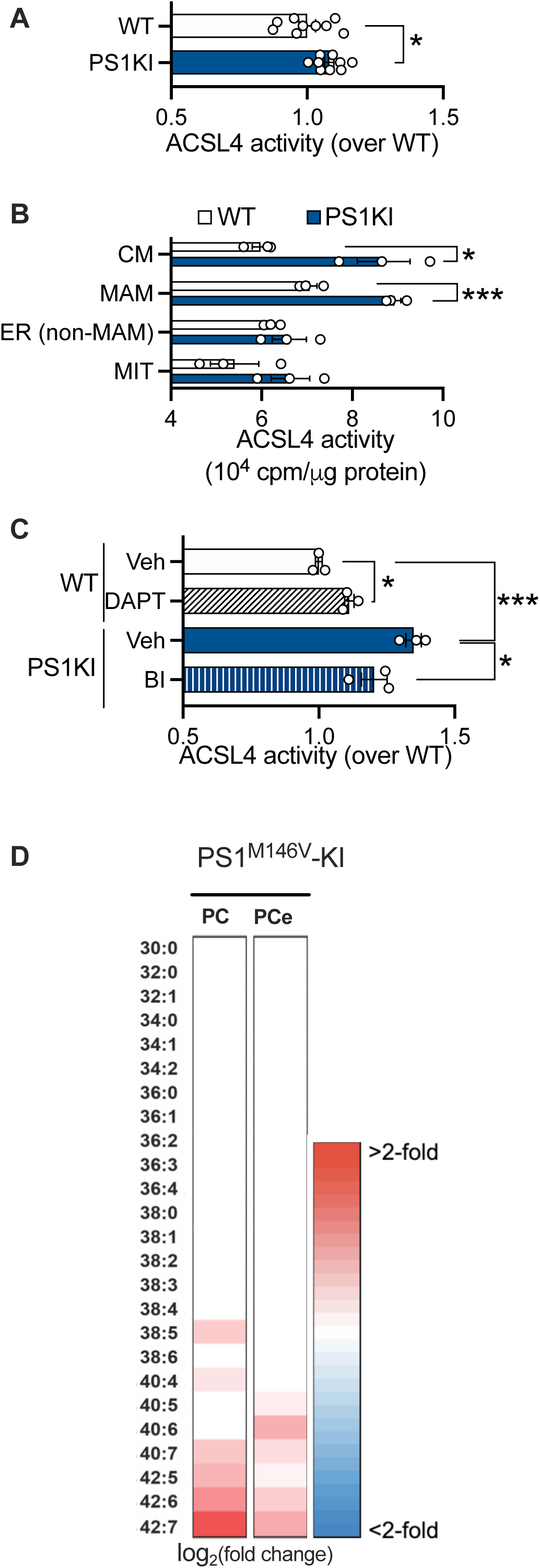
Murine models of AD display ACSL4 upregulation and enrichment of PUFA-containing phosphatidylcholine. **A.** Neonatal cortical neurons from WT or PS1^M146V^-KI mice were cultured and ACSL4 activity was measured. **B.** ACSL4 activity was assayed in membrane preparations of different subcellular fractions obtained from the brains of 1-year old WT and PS1KI mice. CM: crude membranes; MIT: mitochondria; ER: endoplasmic reticulum (non-MAM “bulk” fraction). **C.** ACSL4 activity was measured in WT or PS1^M146V^-KI cultured cortical neurons upon 16h treatment with DAPT or BI, respectively. **D.** Heatmap of fold changes of the levels of PC and PCe species analyzed by lipidomics from 1 year-old PS1^M146V^-KI brains over WT control. Only significant changes are coloured. N=4-5.

In agreement with our previous results, lipidomic analysis of the brain of aged PS1^M146V^-KI mice revealed a significant accumulation of PUFA-containing PC and its ether-linked (PCe) species (**Fig. 4D**). A complementary lipidomic analysis of the P0 brains from the three mouse models found a consistent remodeling of PC, PCe and PE species (**Fig. S3C**).

Consistent with our findings in AD models and supporting translational relevance, fibroblasts derived from both familial and sporadic AD patients exhibited elevated ACSL4 activity compared to age-matched healthy controls (**Fig. 5A**). Lipidomics analysis of these fibroblasts revealed a significant increase in PC and PCe species enriched with PUFAs (**Fig. 5B**), showing a clear increase in the degree of unsaturation of these phospholipids (**Fig. 5C**). Interestingly, a similar lipid signature—characterized by elevated PUFA-containing PC and PCe species—was observed in cortical neurons differentiated from induced-pluripotent stem cells (iPSCs) derived from FAD cases with a pathological mutation in APP (London, V717I) (**Fig. 5D**), and in peripheral blood mononuclear cells (PBMCs) from patients with late-onset AD (**Fig. 5E**). However, this pattern was not mirrored in the plasma from the same individuals, which instead displayed an opposite lipid profile, suggesting cell-type specificity in lipid remodeling associated with AD.

**Figure 5:**
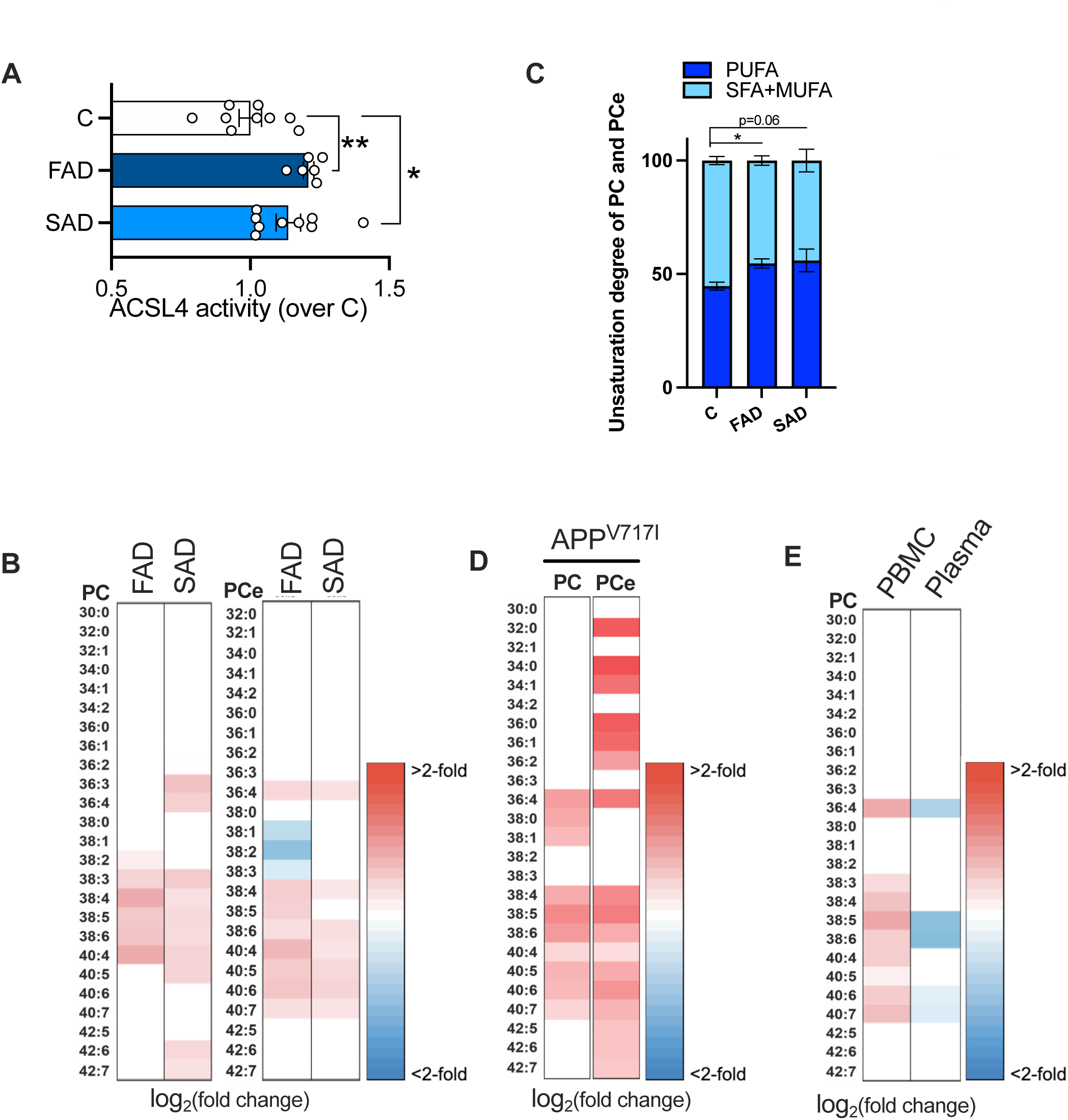
Accumulation of PUFA-containing phosphatidylcholine is a common hallmark in clinical models of AD. **A.** ACSL4 activity was measured in fibroblasts from Familial (FAD) or Sporadic (SAD) AD cases and controls (C). **B.** Heatmap of fold changes of the levels of PC and PCe species analyzed by lipidomics from FAD and SAD fibroblasts over controls. Only significant changes are colored. N=3. **C.** The unsaturation degree of PC and PCe was calculated as the percentage of SFA+MUFA vs PUFA levels. N=3. **D.** Heatmap of fold changes of the levels of PC and PCe species analyzed by lipidomics from APP^V717I^ neurons over controls. Only significant changes are colored. N=4-5. **E.** Lipidomic analysis of PC species obtained from PBMC or plasma from SAD patients over controls. Fold changes are presented. Only significant changes are colored. N=40-50.

Taken together, our data here indicate that C99-mediated increases in MAM and subsequent upregulation of ACSL4 trigger defects in the composition of PC in membranes from AD cells, that likely constitutes an early and critical event in AD progression.

## DISCUSSION

Cellular membranes play a crucial role in compartmentalizing, organizing, and regulating metabolic pathways within the cell. The lipidomic composition of these membranes is a key determinant of cellular function. In particular, the regulation of fatty acid (FA) composition of phospholipids is essential to maintain the biophysical properties of membranes, such as permeability and fluidity. For example, saturated fatty acids (SFA) promote rigidity, whereas polyunsaturated fatty acids (PUFA) enhance fluidity. We show here that the incorporation of polyunsaturated fatty acids (PUFAs), such as arachidonic acid (AA), into phosphatidylcholine (via the Lands cycle) is regulated by the formation of MAM domains and the subsequent activation of FA–metabolizing enzymes such as ACSL4 and LPCAT4. Our findings strongly support the emerging view that MAMs are dynamic functional platforms critical for cellular metabolism and intracellular signaling.

Multiple enzymes contribute to lipid remodeling, however increasing evidence highlights ACSL4 as essential for maintaining PUFA-containing phospholipids that sustain membrane properties, inflammatory mediators, and ferroptosis—a regulated cell death pathway driven by iron-dependent lipid peroxidation [34, 35]. Yet, the regulation of ACSL4 is not fully understood. Our results confirm that ACSL4 expression does not strictly correlate with activity, suggesting additional regulatory mechanisms, including compartmentalization [21]. Notably, we demonstrate here a strong correlation between MAM formation and ACSL4 activation within this domain.

We note that the formation of MAM, as a lipid-raft domain, results from local increases in the levels of cholesterol in the ER delivered from the PM [20]. This change in the lipid environment induces the temporary clustering and conformational modifications of various lipid enzymes, such as ACSL4, that underlie the modulation of their activity. This notion is validated by previous reports showing that alterations in the expression of MFN2, which disrupt the formation of MAM, impact on the regulation of ACSL4, the proportion of PUFA-containing lipids, leukotriene production and susceptibility to ferroptosis [36].

ACSL4 partitioning into specific organelles reflects the broader concept that FA activation is highly compartmentalized. For example, ACSL4 at peroxisomes directs AA toward peroxisomal metabolism, whereas mitochondrial ACSL4 facilitates entry of FAs into β-oxidation. Within MAMs, ACSL4 preferentially channels AA into PC, rather than into other lipid classes (e.g., TAGs, CEs, sphingolipids) [18, 37, 38]. This specificity may reflect localized acyl-CoA pools or a cellular strategy to sequester free FAs, preventing cytosolic accumulation and lipid peroxidation. Importantly, such partitioning may vary among cell and tissues types.

The compartmentalization of ACSL4 in MAM can also modulate its dynamic interaction with other lipid synthesis enzymes, thereby channeling the activated FA into the synthesis or remodeling of different lipid species, directly impacting membrane properties [11]. Our findings show that ACSL4 localization at MAMs promotes functional interaction with LPCAT4, enabling efficient provision of AA-CoA for PC reacylation via the Lands cycle. This remodeling pathway is essential for adjusting membrane composition to cellular demands [9].

Consistent with previous work, ACSL4 and LPCAT3/4 cooperate in PC remodeling with AA, increasing membrane desaturation and sensitivity to ferroptosis [39]. While LPCAT3 and LPCAT4 are functionally similar [40], the preference for ACSL4 interaction with either isoform is currently unknown and may depend on tissue-specific expression or subcellular localization. Also possible, changes in the subcellular localization, phosphorylation state and conformation of ACSL4 can result in different affinities for PUFA-CoAs to be incorporated more readily into different lipids. [41]. Finally, although ACSL4 can activate other PUFAs, such as C22:4 and C22:5 [7, 42], MAM formation may extend beyond ACSL4 to influence other ACSL family members and lipid classes [8, 43]. Despite these complexities, our data support the hypothesis that MAMs promote sequential ACSL4–LPCAT4 reactions, enhancing AA incorporation into PC and thereby altering membrane architecture and signaling microdomains. Dysregulation of this process can impair membrane-dependent functions, contributing to cancer, kidney fibrosis, and metabolic dysfunction-associated steatohepatitis (MASH) [44–47].

In the brain, ACSL4 function is critical for preserving neuronal structural integrity, synaptic homeostasis, and efficient signal transduction [48, 49]. Dysregulated ACSL4 has been linked to neurological disorders, cognitive decline, and neurodegeneration. In particular, in the context of Alzheimer’s disease (AD), ACSL4 is upregulated, disrupting Lands cycle activity and membrane homeostasis [9, 50, 51], while free AA levels have been reported to decline with increasing disease severity [52].

Consistent with this, our data show that AD cells show reduced levels of free AA concomitant with increased AA-containing phospholipids, rendering them vulnerable to lipid peroxidation and inflammatory eicosanoid production [53]. These findings also align with evidence implicating ferroptosis in AD [54], as iron dysregulation may further exacerbate APP expression and Aβ production [55, 56]. However, the mechanism behind these molecular phenotypes and their link to APP metabolism is currently unclear.

Relevant to this, our previous results revealed that the C99 fragment of APP is a potent inducer of MAM formation by promoting cholesterol accumulation in the ER [20]. This positions C99 not as a passive APP byproduct, but as an active regulator of organelle communication and lipid trafficking [58]. Therefore, in AD, we propose that C99 accumulation induces the upregulation of MAM and ACSL4 activation, initiating a pathological feedback loop: increased cholesterol and C99 at the ER promote MAMs, activating ACSL4/LPCAT, disrupting lipid homeostasis, and enhancing APP processing and the concomitant increase in C99 [59].

Altogether, our work demonstrates that MAMs orchestrate a spatially confined lipid circuit, coordinating ACSL4 and LPCAT4 activities to fine-tune lipid remodeling. Under physiological conditions, this ensures adaptive membrane responses to cellular demands. In neurodegenerative disease such as AD, chronic MAM activation disrupts this balance, driving progressive lipid changes detectable in cells and fluids from patients before clinical symptoms appear [43, 60]. These findings establish MAMs as central hubs for lipid homeostasis, linking APP metabolism, ferroptosis, and neurodegeneration. Targeting this MAM-centered metabolic network may thus represent a novel therapeutic strategy for AD and related disorders.

## Materials and Methods

### Cells, mice, PBMCs and plasma

WT and *PSEN1/2* double knock-out (called PS-DKO) mouse embryonic fibroblasts (MEFs) were provided by Dr. Bart De Strooper (KU Leuven, Belgium). APP-WT and *APP/APLP2* double knock-out (called APP-DKO) MEFs were a kind gift from Dr. Huaxi Xu (Sanford-Burnham Medical Research Institute, La Jolla). *MFN2*-knock out cells were a kind gift by Dr. David Chan (California Institute of Technology). AD and control fibroblasts were obtained from the Coriell Institute for Medical Research (Camden, NJ, USA). Other *PSEN1*-mutant FAD cells were kind gifts from Dr. Gary E. Gibson, Dr. Richard Cowburn and Dr. Tom Bird. (see Table S7 for clinical details).

Plasma and peripheral blood mononuclear cells (PBMCs, both derived from the same patient) were obtained from ∼50 sporadic Alzheimer disease (AD) patients and ∼50 age- and sex-matched controls – mainly of Caucasian origin- were provided by Dr. Xiongwei Zhu from the Alzheimer Disease Research Center (ADRC) at Case Western Reserve University (Cleveland, USA).

PS1^M146V^ knock-in and C57BL6 wild-type control mice were obtained from Jackson (#004193) and maintained in the SPF facility at Columbia University Medical Center. 3xTgAD and C57BL6 wild-type controls were bred in the SPF facility at the University of Antioquia. Animals were group-housed (five per cage) in a sound-attenuated room under controlled conditions (12 h light/dark cycle, lights on 07:00–19:00 h; temperature 20 ± 1 °C; humidity 50 ± 10%) with *ad libitum* access to food and water.

Cortical neurons from C57BL6 WT or PS1^M146V^-KI mice were cultured from P0 pups as reported [20]. All mice experiments were performed under the approval of the Institutional Animal Care and Use Committee of the Columbia University Medical Center and the Animal Ethics Committee of the University of Antioquia (Medellín, Colombia).

### Drugs and treatments

Cells were treated with either 5 µM TAPI, 100 nM β-secretase inhibitor IV (BI) or 10 µM DAPT to inhibit α-secretase, β-secretase or γ-secretase, respectively. Neutral sphingomyelinase (SMase) activity was inhibited with 5 µM GW4869. 5µM Triacsin C was used as an ACSL4 inhibitor. 10 µM methyl arachidonyl fluorophosphonate (MAFP) was used to inhibit phospholipase A2. All drugs were incubated for 12–16h and DMSO was used as vehicle.

### Plasmid constructs, RNA silencing and transfections

Cloning of C99^WT^ and C99^MUT^ plasmids was described in [20, 31]. Cells were transfected using Lipofectamine™ 2000 Transfection Reagent in serum-free DMEM for 4-6h following manufacturer’s instructions. For ACSL4 silencing, 25pmol of the Silencer Select s78495 siRNA was transfected using Lipofectamine RNAiMax Reagent. Silencer select siRNA negative control n°2 were used as a control.

### Subcellular fractionation

Purification of ER, crude membranes (CM), mitochondria and MAM was performed and analyzed as described [20].

### Western blotting

After quantification of lysate protein concentrations via BCA assay, equal amounts of protein were boiled in 1xLaemmli buffer and run in Tris-glycine gels. Proteins were detected using the antibodies listed in Reagents Table.

### Immunofluorescence

Mice were deeply anesthetized with ketamine/xylazine and sacrificed. Brains were bisected; the left hemisphere, blood, and cerebrospinal fluid were frozen at –80 °C for biochemical analyses, while the right hemisphere was immersion-fixed in 4% paraformaldehyde (PFA) in 0.1 M phosphate buffer (PB, pH 7.4) at 4 °C for 48 h with daily solution changes. Tissue was cryoprotected in graded sucrose (7%, 25%, 30%), frozen in isopentane, and stored at –80 °C. Coronal sections (50 µm) were prepared using a cryostat (LEICA CM1850 UV) and stored at –20 °C in antifreeze medium (30% ethylene glycol, 30% glycerol in 0.1 M PB). Free-floating sections from the medial bregma were washed in PB, subjected to antigen retrieval (1× citrate buffer, pH 6.0, 15 min, RT), permeabilized (1% Triton X-100, 15 min), and treated to reduce autofluorescence (50 mM NH₄Cl, 15 min; 1% Sudan Black in 70% ethanol, 10 min). Nonspecific binding was blocked with 1% bovine serum albumin (BSA) and 0.3% Triton X-100 in PB for 1 h at RT. Sections were incubated for 72 h at 4 °C with anti-ACSL4 and anti-NeuN in 0.3% BSA/0.3% Triton X-100/PB. After washing, tissues were incubated with Alexa Fluor–conjugated secondary antibodies for 2h at RT. Hoechst was added for nuclear staining during the final 15 min. Sections were washed, mounted with FluorSave, and imaged on an Olympus IX81 epifluorescence microscope with 20× and 60× objectives. Negative controls lacking primary antibodies were included. Images were acquired with a cooled CCD camera and fluorescence intensity was quantified using *ImageJ*.

### PhotoClick cholesterol proteomics in MAM fractions

To study the proteome interacting with cholesterol at MAM, a method developed by [24] was adapted as in [20]. Briefly, cells were incubated in serum-free medium for 2h to remove all exogenous lipids, after which 5 µM PhotoClick cholesterol (Hex-5’-ynyl 3β-hydroxy-6-diazirinyl-5α-cholan-24-oate), previously complexed with an aqueous saturated solution of MβCD (38 mM), was added to the cells and incubated for 4h. Upon washes with DPBS, PhotoClick cholesterol was crosslinked under 365nm-UV (0.75 J/cm^2^, UVC 500 Ultraviolet Crosslinker, Amersham Biosciences), washed again, collected and used for subcellular fractionation as previously described. 500 µg (adjusted to a final 150 µL volume with PBS supplemented with protease inhibitors) of TH and MAM fractions were briefly sonicated and subjected to click chemistry by addition of 500 µM biotin-azide, 100 µM Tris[(1-benzyl-1H-1,2,3-triazol-4-yl)methyl]amine (TBTA), 1 mM CuSO_4_ and 1 mM Tris(2-carboxyethyl)phosphine (TCEP) and incubation for 15 min at room temperature in the dark. Then samples were diluted in 50 mM Tris pH 7.4 with protease inhibitors and incubated overnight under rotation at 4°C with streptavidin beads. Upon several washes with Tris 50 mM pH 7.4, beads were collected by centrifugation at 2,000 rpm for 1min. Samples for proteomic analysis were snap frozen and stored at -80 °C.

### Proteomics analysis

Tandem mass spectrometry after on-bead digestion was performed for proteomic profiling. Proteins were first reduced with 10 mM TCEP and then alkylated with 11 mM iodoacetamide, followed by quenching with 5 mM DTT. Digestion was carried out by incubating the samples with 1 μg of trypsin/LysC mixture overnight at 37°C with shaking at 1400 rpm. The following day, the resulting peptides were transferred to fresh microcentrifuge tubes, and the digestion was terminated by adding 1% (v/v) trifluoroacetic acid. The samples were then centrifuged at 14,000g for 10 minutes at room temperature to remove debris. The supernatants containing digested peptides were desalted and dried using a speed vacuum concentrator. Peptides were reconstituted in 3% acetonitrile with 0.1% formic acid, and 200 ng from each sample was injected for diaPASEF24 analysis using the timsTOF Pro instrument. The raw data from diaPASEF were analyzed against the UniProt Mouse proteome database using the DIA-NN search engine, applying a 1% false discovery rate (FDR) threshold at both the peptide precursor and protein levels. Quantification of the peak areas was performed using the MaxLFQ algorithm, and the resulting data were log2-transformed for further analysis.

Proteomics analysis detected 1647 proteins (Table S1). For functional enrichment analysis, proteins were plugged into the open-source gene ontology analysis site g-profiler for Gene Ontology and KEGG Reactome.

### Measurement of ACSL4 activity

ACSL4 activity was measured as the conversion of ^3^H-arachidonic acid into the soluble ^3^H-arachidonic-CoA. Briefly, 100 µg of membrane fractions were incubated at 37 °C for 10min with 0.1 µCi ^3^H-arachidonic acid in a buffer containing 200mM Tris-HCl pH 7.5, 2.5 mM ATP, 8 mM MgCl2, 2mM EDTA, 20mM NaF, 0.1% Triton X-100. Reactions were initiated by the addition of 0.5mM coenzyme A and terminated by the addition of 3 volumes of isopropyl alcohol, n-heptane and 1M H2SO4 (40:60:1). After addition of 2 volumes of n-heptane, vortexing, and centrifugation at 13000rpm for 5min, the lower aqueous phase (containing soluble ^3^H-arachidonic-CoA) was recovered and subjected to scintillation counting.

### Radioactive fatty acid incorporation assays

0.2 µCi/mL ^3^H-palmitic acid, ^3^H-oleic acid or ^3^H-arachidonic acid were complexed in 2% FA-free BSA and added to the cells for the time periods indicated in serum-free conditions. After extensive washing, cell pellets were harvested. Equal protein amounts were used to extract lipids by using three volumes of chloroform:methanol (2:1 v/v). After vortexing and centrifugation at 8,000 g for 5 min, the organic phase was blown to dryness under nitrogen. Dried lipids were resuspended in 30 µL of chloroform:methanol (2:1 v/v) and applied to a thin layer chromatography plate along with unlabeled standards. A mixture of hexanes/diethyl ether/acetic acid (80:20:1 v/v/v) was used as solvent to separate free FAs, cholesteryl esters (CE), and triacylglycerides (TAG). A two-phase separation was used to separate phosphatidylserine (PS), phosphatidylethanolamine (PE), phosphatidylcholine (PC) and phosphatidylinositol (PI) as previously described [24]. Iodine-stained bands of interest were scraped and counted by scintillation. Percentage of incorporation was measured as the counts in each lipid group divided by the total radioactivity (PS+PE+PC+PI+CE+TAG).

### Deuterated arachidonic acid incorporation and lipidomics analysis

Lipids were extracted from equal amounts of material (30 μg protein/sample). Lipid extracts were prepared via chloroform–methanol extraction, spiked with appropriate internal standards, and analyzed using a 6490 Triple Quadrupole LC/MS system (Agilent Technologies, Santa Clara, CA). Free cholesterol and CE were separated via normal-phase HPLC using an Agilent Zorbax Rx-Sil column (inner diameter 2.1 Å ∼100 mm) under the following conditions: mobile phase A (chloroform:methanol:1 M ammonium hydroxide, 89.9:10:0.1, v/v/v) and mobile phase B (chloroform:methanol:water:ammonium hydroxide, 55:39.9:5:0.1, v/v/v/v); 95% A for 2 min, linear gradient to 30% A over 18 min and held for 3 min, and linear gradient to 95% A over 2 min and held for 6 min. Quantification of lipid species was accomplished using multiple reaction monitoring (MRM) transitions in conjunction with referencing of appropriate internal standards. Values were represented as mole fraction with respect to total lipid (% molarity). For this, lipid mass of any specific lipid is normalized by the total mass of all the lipids measured.

For deuterated arachidonic acid incorporation, 0.2 µM d11-arachidonic acid was complexed in 2% FA-free BSA and added to cells. After extensive washing, lipid extracts were prepared and analyzed as explained above. To detect the transfer of deuterated arachidonic acid (d11-AA) to phospholipids, for negative mode, the fragmentation peak of -314.3 m/z, which corresponds to the negatively charged ion of d11-AA, were scanned. These peaks were cross-checked with the same retention time as the same phospholipid species of normal isotope composition. For positive mode, normal PC peaks were identified first, then shifts by +11.06 m/z at the same retention time with similar fragmentation pattern were examined. Values were later double-confirmed that such identified lipid species with d11-AA followed the same pattern across different experimental groups with normal isotope counterparts.

### Statistical analysis

Data are shown as mean ± SEM. All averages are the result of three or more independent experiments, carried out at different times with different sets of samples. Data distribution was assumed to be normal. The statistical analysis was performed using GraphPad Prism v9.01. Brown-Forsythe test was used to compare variance between groups. Statistical significance was determined by either two-tailed t-test or (repeated measures) two-way ANOVA, followed by Tukey’s or Sidak’s multiple comparison post-hoc test. Welch’s corrections were used for t-test comparison when variance between groups was statistically different. Values of p<0.05 were considered statistically significant. * p<0.05, ** p<0.01, *** p<0.001, **** p<0.0001. The investigators were not blinded when quantifying imaging experiments.

## Data Availability

The mass spectrometry proteomics data have been deposited to the ProteomeXchange Consortium via the PRIDE partner repository with the dataset identifier PXD053282. The lipidomics data is available in Zenodo (#). These data area also provided in the supplemental information/extended data files. All reagents used in this work are commercially available, with manufacturer catalog numbers included in the reagents table. Source data is also provided

## Acknowledgements.

This work was supported by the U.S. National Institutes of Health (R01-AG056387-01 to EA-G; R01NS117538 to EA-S), the National Institute on Aging (1R21AG079574 to GPCG); the Spanish Ministry of Science, Innovation and Universities (PID2021-126818NB-I00 to EA-G; FPI fellowship PRE2022-104771 to AC-F; FPU fellowship 22/00245 to MU; FPU fellowship FPU23/02047 to CA-B); Comunidad de Madrid (2024-T1SAL-GL-31373 Talento César Nombela to JM); the European Union’s Horizon 2020 research and innovation programme under the Marie Skłodowska-Curie grant agreement (MSCA-PF 101106857 to JM); the Minister of Science and Technology Colombia (Minciencias-ICETEX, code #82336 to GPCG); and the CODI-University of Antioquia (to GPCG). We thank Renu Nandakumar for assistance with the lipidomics analysis.

## Author Contributions Statement

Conceived the project: JM and EA-G. Designed experiments: JM and EA-G. Generated experimental data: JM, EA-G, IDS-C, SCA-C, MU, AC-F, CA-B, GPC-G, KRV, EB and NGL. Collected patient samples: XG and FL. Collected/analyzed lipidomics data: FB-A, TDY and EA-G. Wrote the manuscript: JM, GPC-G, EA-G. Critically edited the manuscript: all authors. Approved final version of the manuscript: all authors. We would also like to note that Dr. Lopera, who contributed substantially to this work, passed away during the preparation of the manuscript.

## Competing interests

The authors declare that they have no competing interests. Requests for materials should be addressed to E.A.-G.

---

**Figure S1:**
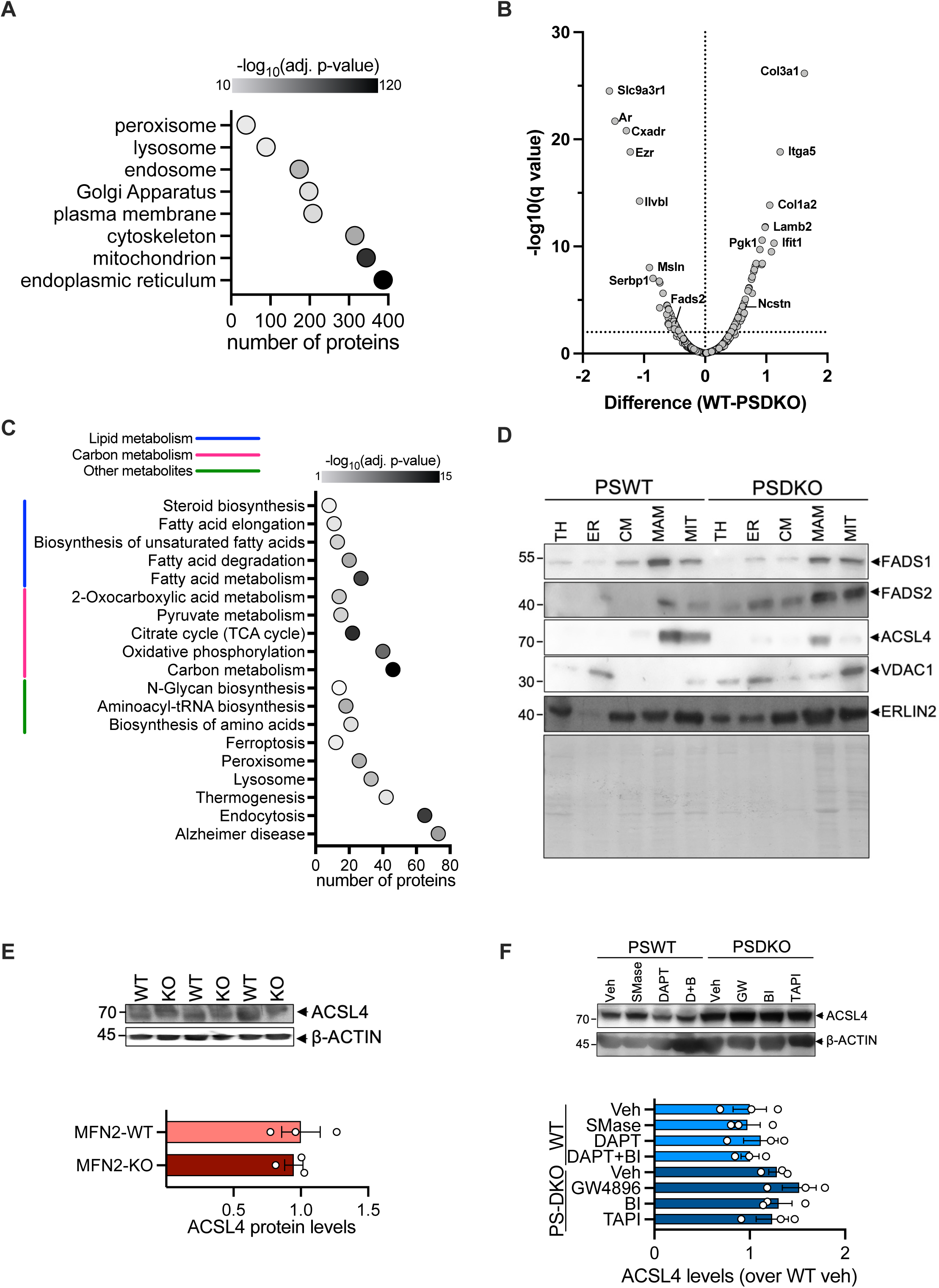
MAM proteomic profiling in response to cholesterol accumulation. **A.** Gene ontology (GO) analysis – cellular component category - from PhotoClick-cholesterol pull-down proteomics. The complete GO analysis for the cellular components is reported in Table S2. **B.** Volcano plot of differentially-abundant proteins detected from PhotoClick-cholesterol pull-downs between WT and PS-DKO MAM fractions. The complete list of proteins detected is included in Table S4. **C.** KEGG pathway enrichment analysis for relevant metabolic pathways. A complete list of categories is found in Table S6. **D.** The levels of FADS1, FADS2 and ACSL4 were assessed in different subcellular fractions obtained from WT or PS-DKO cells. VDAC1 and ERLIN2 were used as mitochondria and MAM enrichment markers, respectively. A Ponceau stain of the membrane is also included. TH: total homogenate, ER: endoplasmic reticulum, CM: crude membranes, MIT: mitochondria. **E.** ACSL4 protein levels measured by western blot in MFN2-KO cells and their WT counterpart. A representative western blot is shown. β-actin was used as a loading control. **F.** ACSL4 protein levels measured by western blot in WT and PS-DKO cells upon treatment with SMase, DAPT, BI, TAPI, GW4896 (GW) or DAPT+BI (D+B), as indicated. A representative western blot is shown. β-actin was used as a loading control.

**Figure S2:**
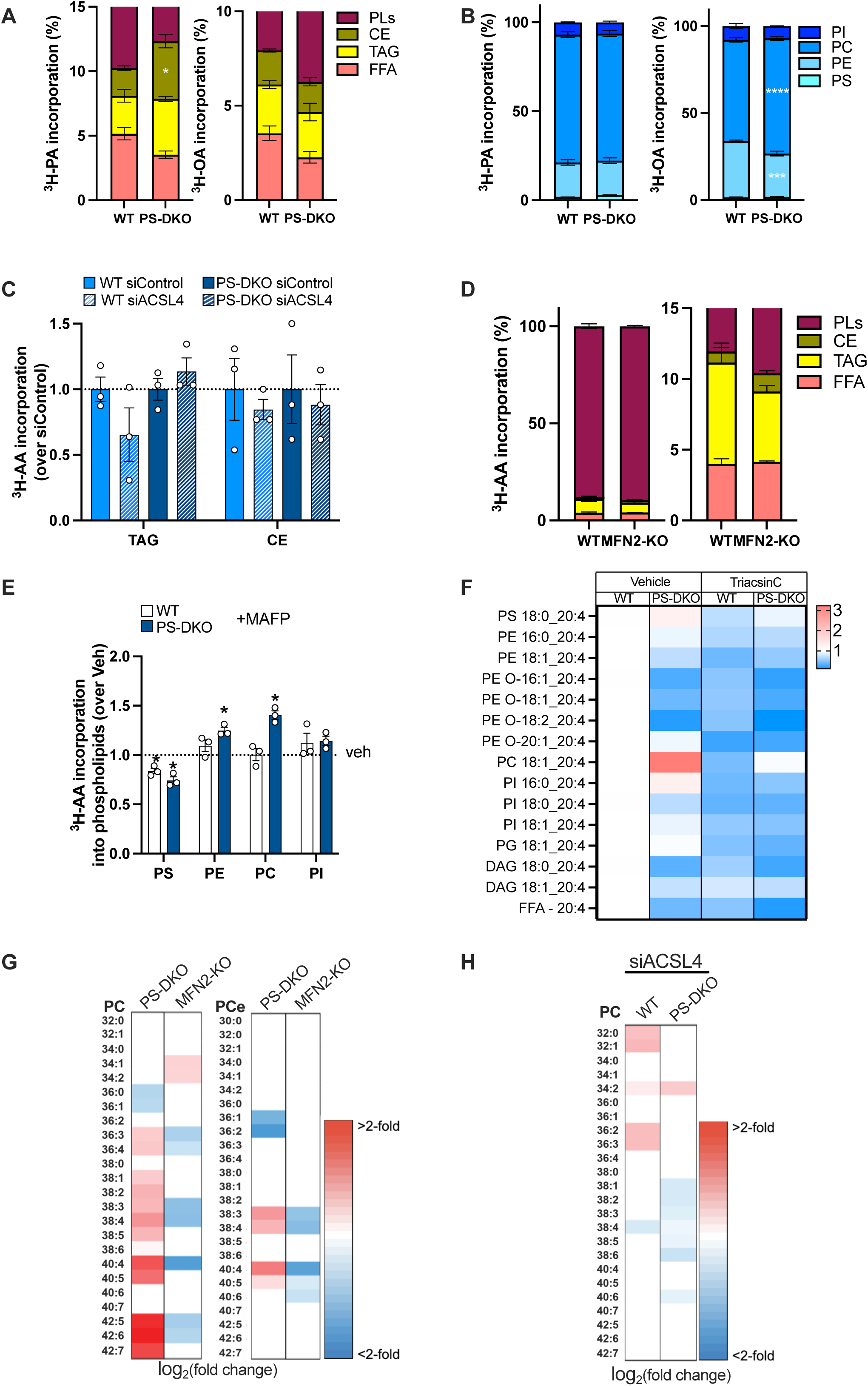
MAM modulation alters the incorporation of fatty acids into different lipid classes. **A.** Comparison of the incorporation of radioactive palmitic acid or oleic acid into specific lipid classes between WT and PS-DKO cells as shown in Fig. 2A-B. **B.** Comparison of the incorporation of radioactive palmitic acid or oleic acid into specific phospholipid classes between WT and PS-DKO cells. **C.** Incorporation of ^3^H-arachidonic acid upon 4h incubation into TAG and CE was assessed in WT and PS-DKO cells silenced for ACSL4. **D.** Incorporation of arachidonic acid into specific lipid classes was assessed in WT and MFN2 - KO cells. **E.** WT and PS-DKO cells were incubated for 4h with radioactive arachidonic acid (^3^H-AA) in the presence of the PLA2 inhibitor MAFP and the incorporation of ^3^H-AA into specific phospholipid classes was measured. Data were normalized over the vehicle-treated conditions (dashed line). **F.** WT and PS-DKO cells were incubated for 4h with deuterated arachidonic acid (dAA) in the presence of the ACSL inhibitor Triacsin C and dAA incorporation was analyzed by lipidomics. Data were normalized over vehicle-treated WT cells. Only significant changes are colored. N=3. **G.** Heatmap of fold changes of the levels of PC and PCe species analyzed by lipidomics from PS-DKO and MFN2-KO MEFs over their control counterparts. Only significant changes are colored. N=3. **H.** Heatmap of fold changes of the levels of PC and PCe species analyzed by lipidomics from WT and PS-DKO cells silenced for ACSL4. Cells transfected with a non-targeting silencer were used as control. Only significant changes are colored. N=3.

**Figure S3:**
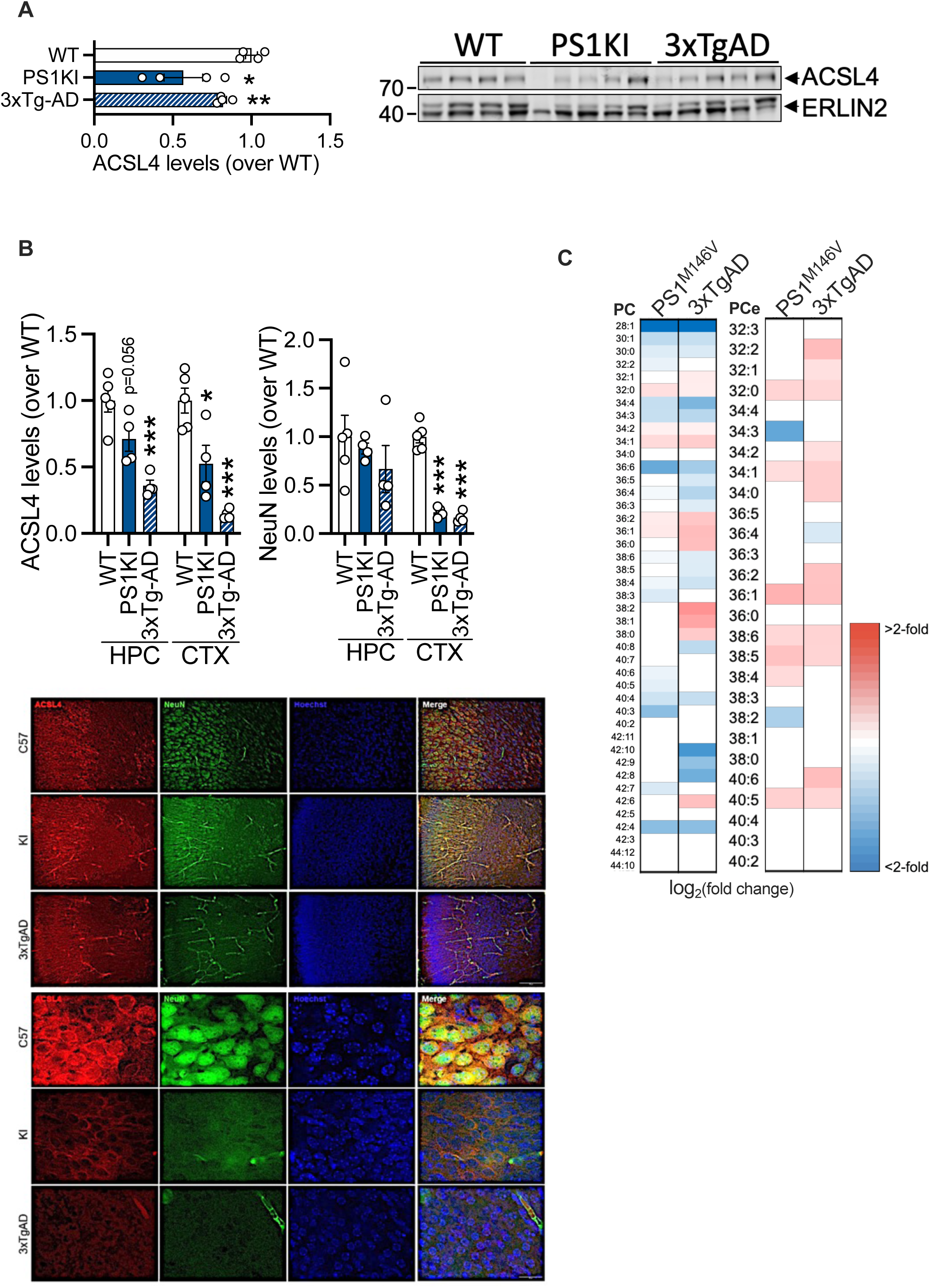
ACSL4 levels and phospholipidomic composition are altered in the hippocampus and the cerebral cortex of PS1^M146V^-KI and 3xTgAD. **A.** Western blotting for ACSL4 in P0 mice. **B.** Triple fluorescence immunostaining for ACSL4 (red), NeuN (green), and Hoechst (blue) at 10X and 60X magnifications. Scale bar= 100, 50 µm. HPC, hippocampus; CTX, cortex. **C.** Heatmap of fold changes with significant p-value for PC and PCe from brains. N=4-5

## Notes

### Competing Interest Statement

The authors have declared no competing interest.

